# The Use of Dipeptide Supplementation as a Means of Mitigating the Negative Effects of Dietary Soybean Meal on Zebrafish *Danio rerio*

**DOI:** 10.1101/2020.04.15.042671

**Authors:** Giovanni S. Molinari, Michal Wojno, Vance J. McCracken, Karolina Kwasek

**Affiliations:** Center for Fisheries, Aquaculture, and Aquatic Sciences, Southern Illinois University, Carbondale, IL, USA; Department of Biological Sciences, Southern Illinois University Edwardsville, Edwardsville, IL, USA

## Abstract

Soybean meal (SBM) is the most common source of protein used to replace fishmeal (FM) in aquaculture diets. SBM inclusion in diets has been found to negatively affect growth and induce intestinal inflammation in fish. The objective of this study was to determine the effect of health-promoting dipeptide (carnosine, anserine, alanyl-glutamine) supplementation into SBM-based feeds on growth performance, intestinal health, and muscle free amino acid composition, an indicator of dietary amino acid availability, in a zebrafish model. There were 5 treatment groups in this study. The first group (**(+) Control**) received a FM-based diet. The second group (**(-) Control**) received SBM-based diet. The last three groups (**Ala-Glu, Car, and Ans**) were fed SBM-based diets, supplemented with alanyl-glutamine, carnosine, and anserine respectively. All groups received their respective diets during 33-59 dph. The Ala-Glu and Car groups experienced a significantly higher weight gain than the (-) Control group, weighing 35.38% and 33.96% more, respectively at the conclusion of the study. There were no significant differences among gene expression between the groups, but Ala-Glu had the highest expression of both nutrient absorption genes measured, PepT1 and *fabp2*. Ala-Glu had significantly longer intestinal villi, and a significantly higher villus length-to-width ratio than the (-) Control group. Among the free amino acid composition, the Car group had a significantly higher post-prandial concentration of lysine, compared to the (-) Control group. The increase in villi surface area and expression of nutrient absorption genes represent an improvement in intestinal absorptive capacity in the Ala-Glu group. The increase in lysine concentration may signify an increase in the retention of protein in the fish in the Car group. The results from this study provide support for the use of alanyl-glutamine and carnosine supplementation as a means of improving growth performance of zebrafish fed with a 100% SBM-based diet.

## Introduction

Many plant protein (PP) sources, including soybean meal (SBM), that are used as alternatives to fishmeal (FM) in fish diets contain anti-nutritional factors such as protease inhibitors, tannins, and saponins, that negatively affect the growth and health performance of both omnivorous [1,2] and carnivorous species [3–5]. These anti-nutritional factors have been found to compromise gut health of fish by inducing intestinal inflammation [1]. Gut health in fish can be assessed on a morphological level by analyzing the status of the intestinal epithelial lining [6–8], or on a molecular level by measuring the expression of pro- and/or anti-inflammatory cytokines within the gut [8–11]. Previously observed signs of SBM-induced intestinal inflammation include shortened mucosal folds, thickened lamina propria, increased number of goblet cells in the epithelium, and wider central stoma within the mucosal folds [3,7,12]. Shortening of mucosal folds within the intestine in particular signify decreased ability to absorb dietary compounds and has also been associated with decreased absorption of both fats and proteins in fish [3]. Genes that have been previously studied to assess intestinal inflammation include pro-inflammatory cytokines interleukin-17A (*il17A*), interleukin-1b (*il1b*), and tumor necrosis factor alpha (*tnf-α*) [10,11]. Marjara et al. [11], found that *il17A* and *il1b* were up-regulated in Atlantic Salmon (*Salmo salar*) intestines afte*r* exposure to SBM. Similar results were observed in zebrafish (*Danio rerio*), where *tnf-α* expression was significantly increased by the inclusion of SBM into the diets [10]. The negative effects of dietary SBM on growth and intestinal health in fish are inevitably limiting its ability to fully replace FM or other high-quality PP concentrates in fish diets. The approach of the present study towards mitigating the negative effects of SBM was focused on the dietary inclusion of health-promoting dipeptides such as carnosine, anserine, and alanyl-glutamine, since their supplementation has been shown to exhibit anti-inflammatory properties in vivo. Two of those dipeptides, carnosine and anserine, are natural dipeptides that originate from meat and contain histidine [13]. Abe [14], found that in pre-adult stages, the level of histidine-containing dipeptides rises in the muscles of fish as body size increases. Consequently, the removal of FM from aquaculture diets means that those dipeptides occurring in fish meat may be lacking in plant-based fish feeds [15].

The effects of carnosine (beta-alanyl-L-histidine) has been extensively researched in mice and rats. Both anti-inflammatory and antioxidative characteristics of carnosine have been reported to be useful for the treatment of chronic alcoholic liver damage in mice [16]. These anti-inflammatory characteristics were also evaluated in the small intestines of rats [17]. Intestines were damaged through irradiation, and mice treated with carnosine through intravenous injection were protected from intestinal inflammation [17]. In fish, there has been a very limited amount of research on the effects of carnosine on intestinal function. One study tested the effects of carnosine supplementation in an all plant-based diet on rainbow trout (*Oncorhyncus mykiss*). Snyder et al. [15] found that carnosine supplementation did not improve cellular muscle development or improve growth. That study, however, utilized a carnosine level of 0.39% in the diet, so it is important to determine if more significant effects would be observed at a higher supplementation level [15]. In addition, the intestinal effects of carnosine supplementation in fish fed plant-based diets have not been researched and, given its success in mammals [17], it is important to determine its viability as a component of plant-based feeds in aquaculture.

Anserine (beta-alanyl-N-methylhistidine) is another dipeptide t found in FM and potentially lacking from plant-based feeds [15]. While the effects of anserine supplementation on intestinal function has yet to be tested in fish, studies have been performed in mammals [18]. For example, anserine has been shown to have antioxidative and anti-glycolytic properties in relation to preservation of blood flow [18]. More research is important in determining whether the antioxidative properties of anserine can support intestinal health of fish fed plant-based feeds.

Alanyl-glutamine has been more extensively studied in the past. In a study using a rat model, alanyl-glutamine supplementation of a standard solution of total parenteral nutrition formula improved intestinal health by protecting mucosal cellularity and function within the intestine [19]. Specifically, increased intestinal villus length was observed in the treatment group that received alanyl-glutamine [19]. Oral supplements of alanyl-glutamine are effective at reducing intestinal oxidative stress and pro-inflammatory responses induced by endotoxemia, the presence of endotoxins in the blood [20]. Protecting mucosal function and reducing oxidative stress in the intestine are important factors in promoting a healthy digestive tract in animals fed PP. Alanyl-glutamine has also been tested on fish species. Positive results on growth and intestinal Na+-K+ ATPase activities were observed [21,22], but those studies did not involve the prior induction of intestinal inflammation or other damage compromising the functional capacity of the intestine.

Research involving the supplementation of the dipeptides gives important insight into the use of dipeptide supplementation as a means of reducing the negative effects of SBM-induced intestinal inflammation. However, as the majority of the previous studies were done on mammals, it is important to assess whether the health-promoting properties of these dipeptides are transferrable to fish species. Another unknown regarding supplementation of dipeptides is whether anti-oxidative and anti-inflammatory properties of the dipeptides are enough to protect the intestine and improve feed utilization in fish fed a PP-based diet. Therefore, the objective of this study was to determine the effect of supplementation of PP-based feeds with dipeptides (carnosine, anserine, alanyl-glutamine) on growth performance, any morphological and/or molecular changes associated with the intestinal lining, as well as muscle free amino composition used as an indicator of dietary amino acid availability, in the zebrafish model. We hypothesized that the dipeptides reduce intestinal inflammation and the fish fed PP diets supplemented with the tested dipeptides will experience more efficient feed utilization and hence, increased growth.

## Materials and methods

### Animal husbandry

This study utilized zebrafish as a model organism. Zebrafish can feed on both plant and animal protein sources, representing nutritional pathways for both carnivores and omnivores [23]. In addition, zebrafish serves as a good model to analyze diet-induced intestinal inflammation in aquaculture species [24]. The feeding trial was conducted in the Center for Fisheries, Aquaculture, and Aquatic Sciences at Southern Illinois University-Carbondale (SIUC), IL. All experiments were carried out in strict accordance with the recommendations in the Guide for the Care and Use of Laboratory Animals of SIUC. The SIUC Institutional Animal Care and Use Committee approved all of the protocols performed. All researchers were trained in accordance with SIUC Institutional Animal Care and Use Committee requirements. During fish handling, anesthesia was performed using water bath immersion in tricaine methanesulfonate (MS222) at a recommended concentration (0.16 mg/ml), and all efforts were made to minimize pain, stress, and discomfort in the animals. Humane endpoints were used in this study; fish were euthanized within 24 hours if they showed significant signs of deteriorating quality of life. Fish were observed each day for signs of distress. No fish in this study had to be euthanized prior to the completion of the experiment.

All experiments were carried using a recirculated aquaculture system (Pentair Aquatic Eco-systems, Cary, NC). The system consisted of 3 L and 5 L tanks, stacked in rows. Water in the system was constantly filtered and recirculated using a mat filter, UV light, a carbon filter, and biofiltration. The average water temperature was 27.1 ± 0.2°C, the pH was 7.01 ± 0.23, and the salinity was kept between 1 and 3 ppt (higher during the first feeding to prolong the viability of the live food [25]. The photoperiod consisted of 14 hours of darkness and 10 hours of light, with the overhead illumination from 08:00-18:00.

### Feed preparation and formulation

Dry components of feeds were ground to a fine particle size (0.5-0.25 mm) using a centrifugal mill (Retsch 2M 100, Haan, Germany). Once ground, components were mixed (Farberware Mixer, Fairfield, CA) to achieve uniform dispersion of all ingredients within the mix. After mixing, feeds were forced through an extruder (Caleva Extruder 20, Sturminster Newton Dorset, England) and then spheronized (Caleva Multibowl Spheronizer, Sturminster Newton Dorset, England) to obtain solid, spherical pellets. All feeds were then freeze-dried (Labconco FreeZone 6, Kansas City, MO) to remove moisture. Pellets were separated by size using a vibratory sieve shaker (Retsch AS 200 Basic, Haan, Germany) to appropriate sizes. All diets were isoenergetic and isonitrogenous. Crude protein level of the diets was 48.9%, and the crude lipid level was 15%. The SBM diet replaced 100% of FM with a combination of SBM and soy protein concentrate. Soy protein concentrate was included as necessary to adjust dietary crude protein while leaving room for other ingredients in the formulation, including a minimum level of starch to allow expansion of the experimental diets. Feed formulations for this study are listed in Table 1. The SBM diet served as the PP-based diet. Dipeptides were dissolved in water and added to their respective diets after mixing of the dry ingredients. Carnosine was supplemented at a 1% level within the diet, a level 2.5 times the level previously tested in fish [15]. Alanyl-glutamine was also supplemented at a 1% level within the diet, matching the level that has been tested on fish species in past studies [26,27]. Anserine was supplemented into the diet at a level of 0.035%, 10 times the concentration found in fish muscle.

**Table 1.** Feed formulation of experimental diets

| **Ingredients (g/100g)** | **FM** | **SBM** | **Carnosine** | **Ala-Glu** | **Anserine** |
| --- | --- | --- | --- | --- | --- |
| Fishmeal<sup>1</sup> | 63.7 | 0 | 0.0 | 0.0 | 0.0 |
| Soybean Meal<sup>2</sup> | 0.0 | 45.5 | 45.5 | 45.5 | 45.5 |
| Soy Protein Concentrate<sup>3</sup> | 0.0 | 16.0 | 14.9 | 14.9 | 16.0 |
| Krill Meal<sup>4</sup> | 10.0 | 10.0 | 10.0 | 10.0 | 10.0 |
| CPSP<sup>5</sup> | 5.0 | 5.0 | 5.0 | 5.0 | 5.0 |
| Dextrin<sup>3</sup> | 5.4 | 0.0 | 0.0 | 0.0 | 0.0 |
| Fish Oil<sup>6</sup> | 4.2 | 7.8 | 7.8 | 7.8 | 7.8 |
| Soy Lecithin<sup>6</sup> | 5.0 | 5.0 | 5.0 | 5.0 | 5.0 |
| Mineral Mix<sup>3</sup> | 2.5 | 2.5 | 2.5 | 2.5 | 2.5 |
| CaHPO<sub>4</sub><sup>7</sup> | 0.0 | 1.5 | 1.5 | 1.5 | 1.5 |
| Vitamin Mix<sup>3</sup> | 2.0 | 2.0 | 2.0 | 2.0 | 2.0 |
| Vitamin C<sup>8</sup> | 0.1 | 0.1 | 0.1 | 0.1 | 0.1 |
| Choline Chloride<sup>3</sup> | 0.1 | 0.1 | 0.1 | 0.1 | 0.1 |
| Methionine<sup>3</sup> | 0.0 | 0.5 | 0.5 | 0.5 | 0.5 |
| Lysine<sup>3</sup> | 0.0 | 2.0 | 2.0 | 2.0 | 2.0 |
| Threonine<sup>3</sup> | 0.0 | 0.1 | 0.1 | 0.1 | 0.1 |
| Taurine<sup>3</sup> | 1.0 | 1.0 | 1.0 | 1.0 | 1.0 |
| Carnosine<sup>9</sup> | 0.0 | 0.0 | 1.0 | 0.0 | 0.0 |
| Ala-Glu<sup>9</sup> | 0.0 | 0.0 | 0.0 | 1.0 | 0.0 |
| Anserine<sup>9</sup> | 0.0 | 0.0 | 0.0 | 0.0 | 0.035 |
| Guar Gum<sup>3</sup> | 1.0 | 1.0 | 1.0 | 1.0 | 1.0 |

FM – fishmeal, SBM – soybean meal, CPSP – fish protein hydrolysate, Ala-Glu – alanyl-glutamine
<sup>1</sup>Omega Protein, Reedville, VA, USA
<sup>2</sup>Premium Feeds, Perryville, MO, USA
<sup>3</sup>Dyets Inc, Bethlehem, PA, USA
<sup>4</sup>Florida Aqua Farms, Dade City, FL
<sup>5</sup>Soluble fish protein concentrate, Sopropêche, France
<sup>6</sup>MP Biomedicals, Solon, OH, USA
<sup>7</sup>Acros Organics, NJ, USA
<sup>8</sup>Argent Aquaculture, Redmond, WA, USA

### Experimental design and feeding regimen

At 33 dph zebrafish, weighing 0.031g (± 0.001) were randomly distributed into 15 (3L) tanks, with 50 fish in each tank. There were 5 treatment groups in this study. The first group (**(+) Control**) received a FM-based diet. The second group (**(-) Control**) received SBM diet. The remaining groups received SBM-based diets supplemented with either carnosine (**Car**), anserine (**Ans**) or alanyl-glutamine (**Ala-Glu**). All groups received their respective diets during 33-59 dph. The fish were fed with a restricted feeding rate of 8% of the total biomass per day.

### Sampling and measuring

At the conclusion of the study, three fish from each tank were sampled and stored in a 10% formalin solution for histology analyses, and 3 fish in each tank were euthanized with MS-222 and intestines were harvested and stored in RNALater (Sigma-Aldrich, St. Louis, MO, USA) for future analysis of gene expression and assessment of intestinal health. For free amino acid (FAA) analysis, three fish were sampled from each tank at 3 and 24 hours after feeding at the conclusion of the study and stored at −80 oC until analysis.

Fish were weighed on a weekly basis beginning at 33 dph in order to measure biomass and adjust feeding rates. The average weight per fish of each group was measured at the end of the study, as was the percent weight gain of each group during the study. Average weight was calculated for each tank by dividing the final biomass by the number of fish in the tank. The data shown for each group are average weight per fish for the 3 replicates of the group. Weight gain was calculated by subtracting the initial weight (at 33 dph) of the tank from the total weight at the conclusion of the study (59 dph). Weight gain was then divided by initial weight and multiplied by 100 to obtain weight gain percentage. Survival was assessed for the duration of the study (33-59 dph) by dividing the final number of fish in the tank by the initial number of fish (50) and multiplying that number by 100.

### Gene expression analysis

Intestinal samples were processed using TRIzol Reagent (Ambion, Foster City, CA, USA) and RNA was extracted and purified using the On-Column PureLink™ DNase Treatment (PureLink™ RNA Mini Kit and PureLink™ DNase, Invitrogen, Carlsbad, CA, USA) following the manufacturer’s instructions. Once purified, the nanograms/µl of each RNA sample was obtained using a spectrophotometer (Nanodrop 2000c, Thermo Fisher Scientific, Waltham, MA, USA). From this point, 2 µg of RNA from each sample was reverse transcribed using the High Capacity cDNA Reverse Transcription Kit (Applied Biosystems, Foster City, CA, USA) to obtain a 20 µl cDNA solution. The cDNA solutions were then added to a tube with 380 µl of water, to produce the cDNA sample for each tank. Gene expression of each cDNA sample was measured using a Bio-Rad (Hercules, CA, USA) CFX96™ Real-Time PCR System. Each qPCR reaction mixture (20 µl) contained 9 µl of cDNA sample, 10 µl of PowerUp™ SYBR™ Green Master Mix (Thermo Fisher Scientific, Waltham, MA, USA), and 0.5 µl of 800 nMol each of forward and reverse primers. Primers (Table 2) were synthesized by Integrated DNA Technologies (Coralville, IA, USA). Each qPCR reaction was run in technical duplicates. The qPCR cycle consisted of 95oC for 10 min, followed by 40 cycles of 95oC for 20 seconds and 60oC for 35 seconds, followed by a melting curve to ensure the amplification of only a single product in each well. Relative gene expression was calculated using the 2∆∆Ct method, normalizing the target gene expression to the expression of *ef1a* (reference gene) [10], and comparing the expression to that of the (-) Control group.

**Table 2.**
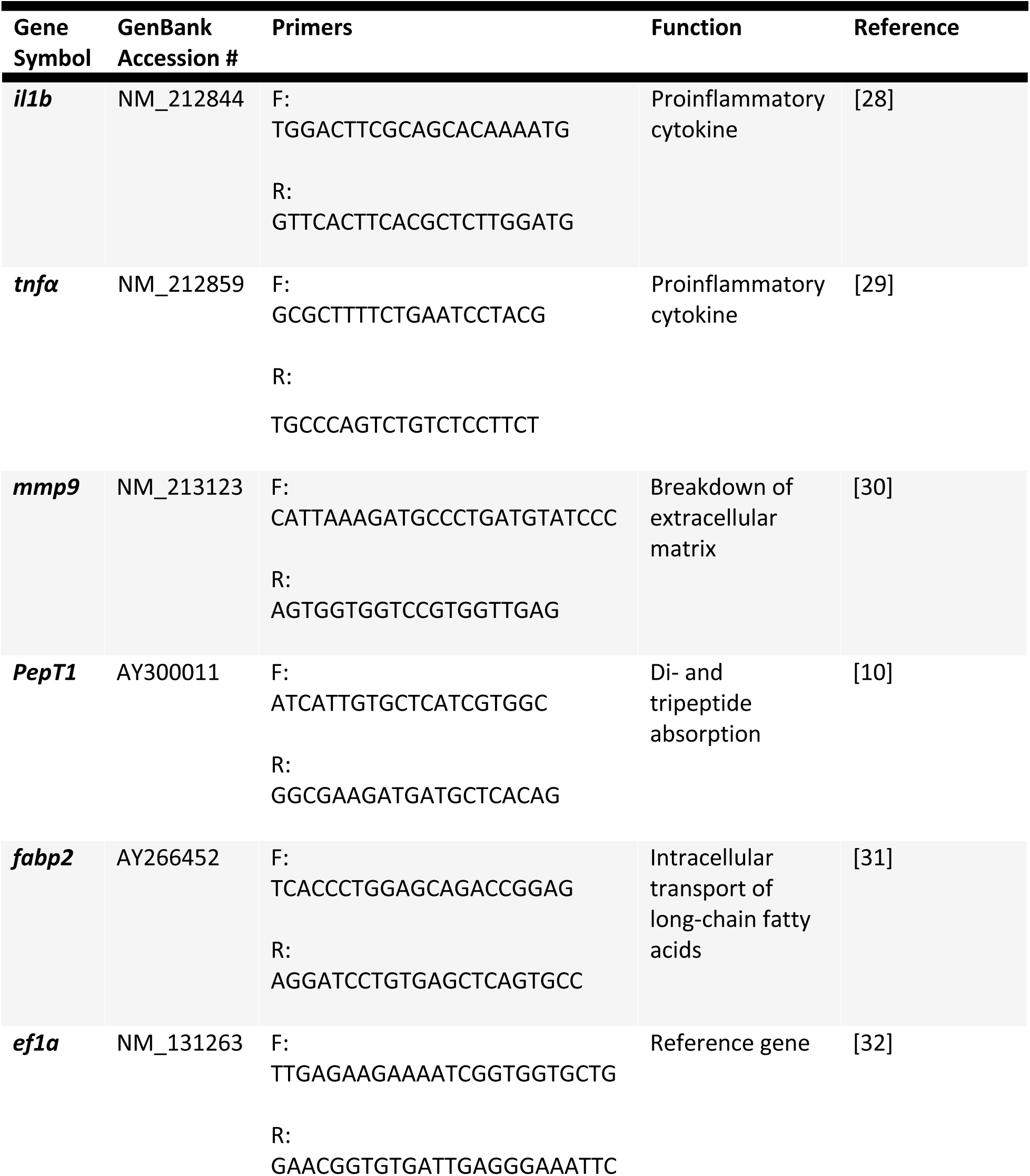
Primers used for gene expression analysis

### Histological analysis

Histological slides were prepared by Saffron Scientific Histology Services (Carbondale, IL). Intestines previously fixed in 10% neutral buffered formalin were processed to paraffin using a Sakura enclosed automated tissue processor (Netherlands). The three representative areas of zebrafish intestines were oriented for cross sections embedded together in the same block. Five-micrometer serial sections were cut with a Leica manual microtome (Buffalo Grover, IL) and placed on water bath at 44oC. Sections were placed on positive charged slides. After drying, slides were stained with hematoxylin and eosin and cover-slipped using acrylic mounting media. The histological analysis focused on the hind-gut portions of fish digestive tract. Pictures of each slide at 100X magnification were obtained using a microscope (Leica DMI 300B) and camera (Leica DMC 290) combination, with the software LAS V4.4 (Leica Camera, Wetzler, Germany). From these pictures, individual lengths and widths were taken of intact villi using ImageJ (NIH, Betheseda, MD, USA). Length and width data were measured in the proximal portion of the intestine. Villus length was measured from the tip of the villus to the luminal surface, and villi width was measured across the base of the villus at the luminal surface. The length-to-width ratio of each villus was determined by dividing the length by the width.

### Free amino acid analysis

Heads, caudal fins and dorsal parts of frozen fish (−80ºC) were removed and the remaining part (muscle) was used for the analysis. Muscle samples of three fish from each tank were combined and homogenized together with 0.1 mol/l HCl in 1:9 (w/v) and spun at 12 000 g (4ºC, 15 min). Supernatants were collected, filtered (Milipore, 10 kDa cutoff at 15 000 g, 4ºC, 30 min) and later diluted with 0.1 mol/l HCl (1:19 v/v) containing norvaline and sarcosine (40 µmol/l) as internal standards. Blanks (0.1 M HCl + 40 μmol/l norvaline and sarcosine) and external standards (Sigma acid/neutral and basic amino acids) were prepared along with the sample preparation. The same concentration of glutamine in 0.1 M HCl as an external standard was prepared and added to the basic amino acids standard. Free amino acids were quantified using Shimadzu Prominence Nexera - i LC-2040C Plus (Shimadu, Japan) according to the Shimadzu protocol No. L529 with modifications. Free amino acid concentrations (expressed as µmol/kg wet body weight) were calculated in LabSolutions software version 5.92 (Shimadzu, Japan) using internal and external standards.

### Statistical analysis

Results are presented as means (± standard deviation). One-way ANOVA was used to test the data, and an LSD test was run to test the differences between groups. Differences between groups are considered significant at *p* values < 0.05. Statistical analysis was performed using R software.

## Results

### Growth performance

The supplementation of alanyl-glutamine and carnosine had a significant effect on the growth performance of zebrafish (Table 3). Both Ala-Glu and Car groups had a significantly higher average weight at the conclusion of the study, compared to the (-) Control group and not different with (+) Control. Both groups also had a significantly higher weight gain during the study, compared to the (-) Control group and not different with (+) Control. The Ans group did not experience a significant improvement in growth performance. There was no difference in survival among the groups, as no mortalities occurred during the study.

**Table 3.**
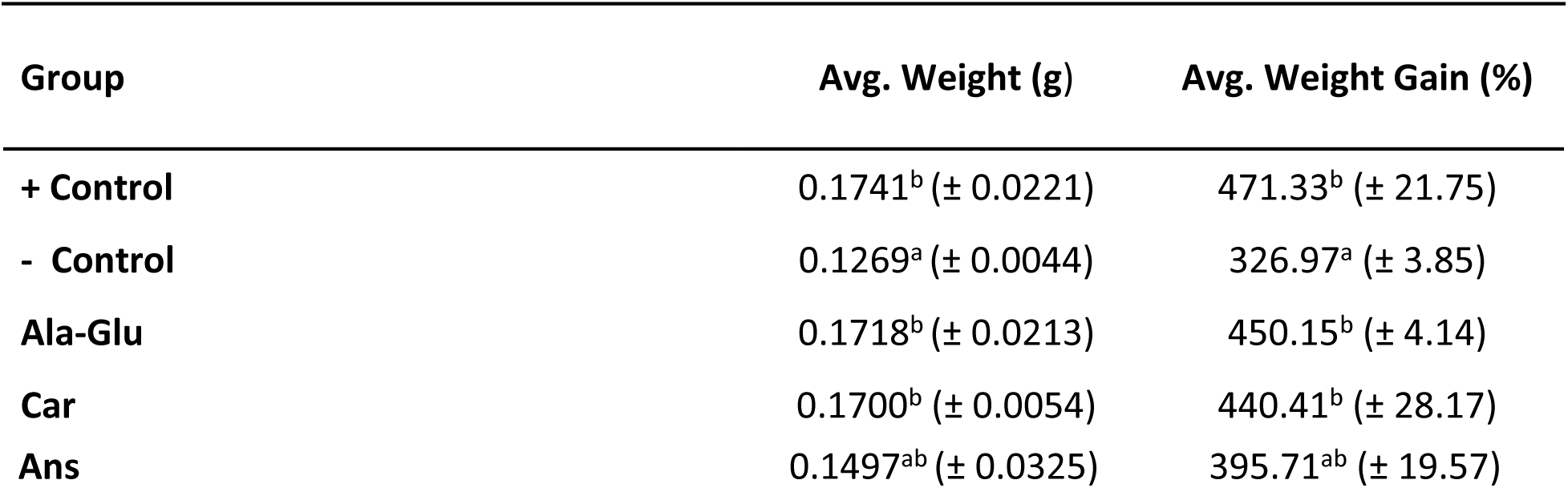
Treatment effect on growth performance measures. Values are presented as means (± std. dev). Superscript letters indicate statistical significance between groups. The significance was determined using a One-Way ANOVA and a LSD Test with a p value <0.05.

### Intestinal health

There were no significant differences in intestinal expression of *il1b*, *il10*, *tnfα*, *mmp9*, *fabp2*, or PepT1 among the groups (Figure 1). One trend observed in the data is that both nutrient absorption genes, PepT1 and *fabp2* tended to be numerically higher in Ala-Glu group compared to (-) Control, Car and Ans groups.

**Fig 1.**
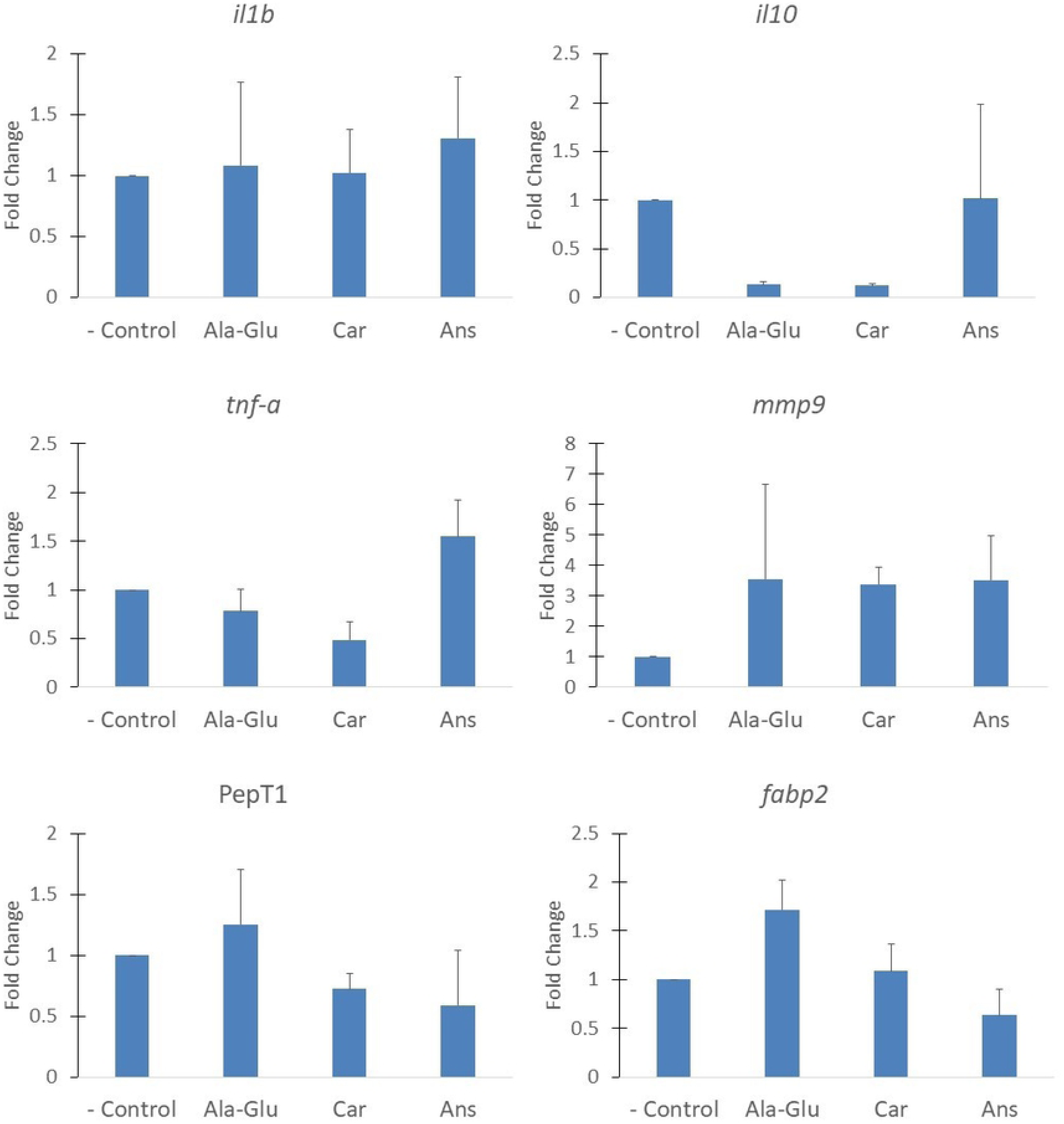
Intestinal gene expression. Relative gene expression of the group is represented as a fold change relative to the negative control group (fold change=1). Values provided are mean fold change + S.E.M (standard error of the mean). Results of One-way ANOVA and LSD test are shown on graphs when significant. Significant differences (p<0.05) between groups are represented by different letters.

The results for intestinal histology are presented in Table 4. Ala-Glu and Ans had significantly longer proximal villi than the (-) Control and Car groups. There were no significant differences in the widths of the intestinal villi between groups. Using the length and width measurements of the intestinal villi, a length-to-width ratio for the villi was calculated to represent the surface area available for nutrient absorption. The Ala-Glu group had a significantly higher length-to-width ratio of proximal villi than both the (-) Control and Car groups. The Ans group had a significantly higher length-to-width ratio than the Car group, and the length-to-width ratio of the (+) Control group was not significantly different from any of the experimental groups.

**Table 4.**
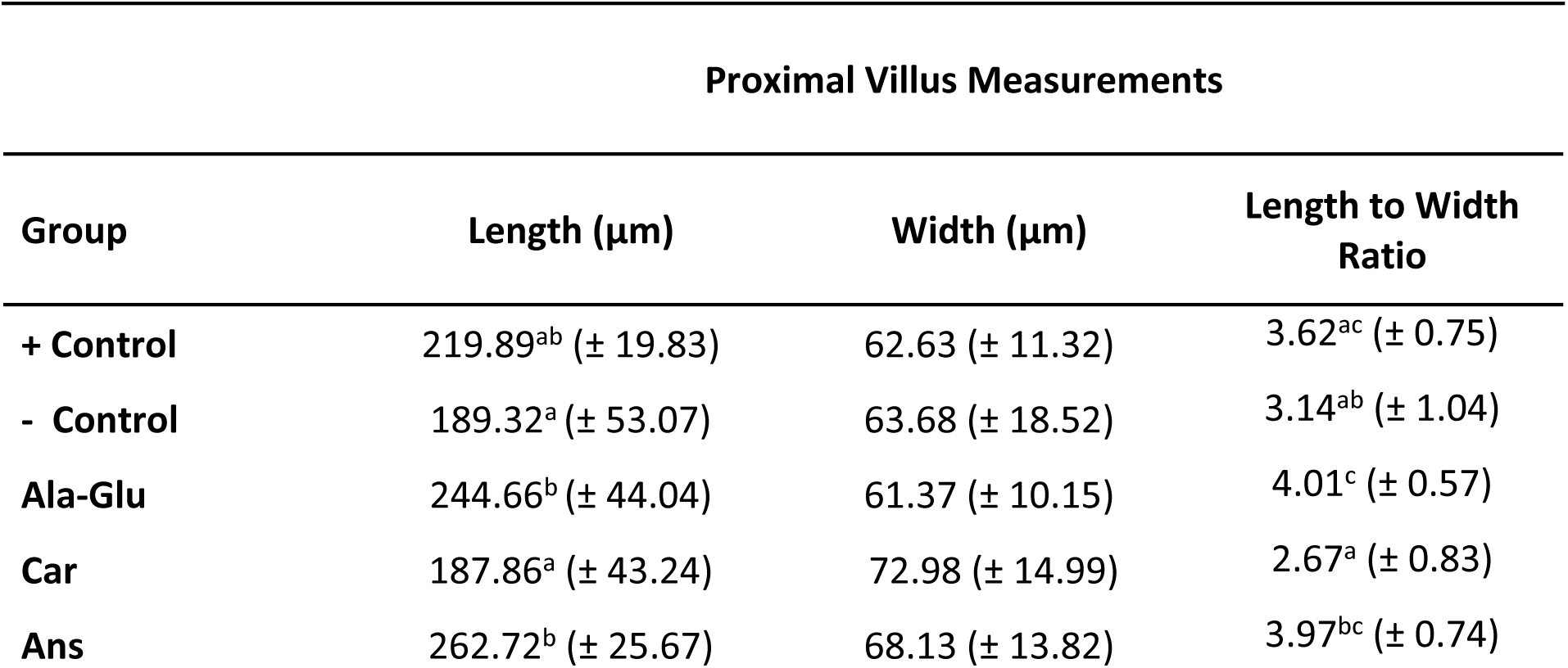
Treatment effect on the villus measurements in the proximal portion of the intestine. Values are presented as means (± std. dev). Superscript letters indicate statistical significance between groups. The significance was determined using a One-Way ANOVA and an LSD test with a p value <0.05.

| Proximal Villus Measurements |  |  |  |
| --- | --- | --- | --- |
| Group | Length (μm) | Width (μm) | Length to Width Ratio |
| + Control | 219.89<sup>ab</sup> (± 19.83) | 62.63 (± 11.32) | 3.62<sup>ac</sup> (± 0.75) |
| - Control | 189.32<sup>a</sup> (± 53.07) | 63.68 (± 18.52) | 3.14<sup>ab</sup> (± 1.04) |
| Ala-Glu | 244.66<sup>b</sup> (± 44.04) | 61.37 (± 10.15) | 4.01<sup>c</sup> (± 0.57) |
| Car | 187.86<sup>a</sup> (± 43.24) | 72.98 (± 14.99) | 2.67<sup>a</sup> (± 0.83) |
| Ans | 262.72<sup>b</sup> (± 25.67) | 68.13 (± 13.82) | 3.97<sup>bc</sup> (± 0.74) |

### Free Amino Acids

The results for muscle FAA postprandial levels 3-hours after feeding are presented in Table 5. Out of 20 FAA analyzed, only 9 showed statistical differences among groups. Among the indispensable free amino acids (IDAA), the concentration of histidine was significantly higher in the Ans group compared to the (+) Control and Car group (Figure 2). However, none of the supplemented groups had a significantly different concentration of histidine compared to the (-) Control group. Phenylalanine was significantly higher in all groups except Ans, compared to the (+) Control group. The concentration of lysine was significantly higher in Car compared to both of the control groups. The concentration of tryptophan was significantly higher in the Car group compared to the (+) Control group, but there were no significant differences among the rest of the groups (Figure 2).

**Table 5.** Dietary treatment effect on the muscle concentration of free amino acids. The values presented are post-prandial levels, 3 hours after feeding. Values are presented as means (± std. dev). Superscript letters indicate statistical significance between groups. Significance was determined using a One-Way ANOVA and an LSD Test with a p value <0.05. Indispensable amino acids (IDAA) = Ile, Leu, Lys, Met, Phe, Thr, Trp, Val, Arg, and His. Dispensable amino acids (DAA) = Ala, Asp, Asn, Glu, Gln, Gly, Pro, Ser, and Tyr. TFAA = Total Free Amino Acids.

| Amino Acid ($\mu\text{mol/g}$) | + Control | - Control | Ala-Glu | Car | Ans |
| --- | --- | --- | --- | --- | --- |
| Aspartic Acid | 0.38 (± 0.16) | 0.24 (± 0.03) | 0.36 (± 0.17) | 0.27 (± 0.05) | 0.33 (± 0.04) |
| Glutamic Acid | 1.59 (± 0.39) | 1.57 (± 0.06) | 1.64 (± 0.27) | 1.92 (± 0.46) | 1.70 (± 0.21) |
| Asparagine | 0.39<sup>a</sup> (± 0.13) | 0.83<sup>b</sup> (± 0.28) | 1.10<sup>b</sup> (± 0.23) | 0.86<sup>b</sup> (± 0.29) | 0.91<sup>b</sup> (± 0.06) |
| Serine | 0.55 (± 0.22) | 0.74 (± 0.59) | 0.82 (± 0.19) | 0.72 (± 0.40) | 0.59 (± 0.09) |
| Glutamine | 1.29 (± 0.38) | 1.36 (± 0.28) | 1.46 (± 0.19) | 1.38 (± 0.22) | 1.31 (± 0.02) |
| Histidine | 7.82<sup>a</sup> (± 1.06) | 9.84<sup>bc</sup> (± 1.59) | 9.51<sup>ac</sup> (± 1.47) | 8.92<sup>ab</sup> (± 0.53) | 11.07<sup>c</sup> (± 0.22) |
| Glycine | 3.39<sup>c</sup> (± 0.97) | 1.70<sup>b</sup> (± 0.59) | 0.83<sup>ab</sup> (± 0.14) | 0.83<sup>ab</sup> (± 0.19) | 0.63<sup>a</sup> (± 0.11) |
| Threonine | 1.27 (± 0.31) | 1.05 (± 0.30) | 0.68 (± 0.03) | 0.64 (± 0.04) | 0.96 (± 0.27) |
| Arginine | 0.40<sup>a</sup> (± 0.21) | 0.67<sup>ab</sup> (± 0.24) | 0.73<sup>ab</sup> (± 0.18) | 0.81<sup>b</sup> (± 0.28) | 0.59<sup>ab</sup> (± 0.18) |
| Alanine | 0.80<sup>b</sup> (± 0.10) | 0.63<sup>a</sup> (± 0.07) | 0.54<sup>a</sup> (± 0.04) | 0.52<sup>a</sup> (± 0.04) | 0.52<sup>a</sup> (± 0.06) |
| Tyrosine | 0.07<sup>a</sup> (± 0.01) | 0.11<sup>b</sup> (± 0.03) | 0.11<sup>b</sup> (± 0.01) | 0.11<sup>b</sup> (± 0.02) | 0.08<sup>ab</sup> (± 0.00) |
| Methionine | 2.38 (± 0.28) | 2.37 (± 0.19) | 2.10 (± 0.01) | 2.25 (± 0.22) | 2.16 (± 0.11) |
| Valine | 0.71 (± 0.13) | 0.66 (± 0.04) | 0.65 (± 0.02) | 0.72 (± 0.12) | 0.63 (± 0.04) |
| Tryptophan | 0.02<sup>a</sup> (± 0.00) | 0.03<sup>ab</sup> (± 0.01) | 0.02<sup>ab</sup> (± 0.00) | 0.03<sup>b</sup> (± 0.01) | 0.02<sup>ab</sup> (± 0.01) |
| Phenylalanine | 0.07<sup>a</sup> (± 0.02) | 0.19<sup>b</sup> (± 0.05) | 0.18<sup>b</sup> (± 0.03) | 0.22<sup>b</sup> (± 0.10) | 0.16<sup>ab</sup> (± 0.01) |
| Isoleucine | 0.21 (± 0.09) | 0.22 (± 0.06) | 0.26 (± 0.02) | 0.30 (± 0.13) | 0.19 (± 0.08) |
| Leucine | 0.36 (± 0.15) | 0.38 (± 0.11) | 0.43 (± 0.04) | 0.51 (± 0.23) | 0.32 (± 0.11) |
| Proline | 2.26 (± 0.58) | 1.67 (± 0.79) | 2.77 (± 0.56) | 2.45 (± 0.65) | 2.60 (± 0.54) |
| Lysine | 1.17<sup>a</sup> (± 0.51) | 1.16<sup>a</sup> (± 0.45) | 1.80<sup>ab</sup> (± 0.20) | 2.10<sup>b</sup> (± 0.53) | 1.62<sup>ab</sup> (± 0.54) |
| **IDAA** | 14.40 (± 2.08) | 16.57 (± 2.49) | 16.89 (± 2.01) | 16.95 (± 2.34) | 17.91 (± 0.99) |
| **DAA** | 10.73<sup>b</sup> (± 0.83) | 8.85<sup>a</sup> (± 1.26) | 9.63<sup>ab</sup> (± 1.12) | 9.05<sup>ab</sup> (± 0.91) | 8.67<sup>a</sup> (± 0.24) |
| **TFAA** | 25.13 (± 2.93) | 25.42 (± 3.67) | 26.52 (± 3.13) | 26.00 (± 3.08) | 26.58 (± 0.87) |

**Fig 2.**
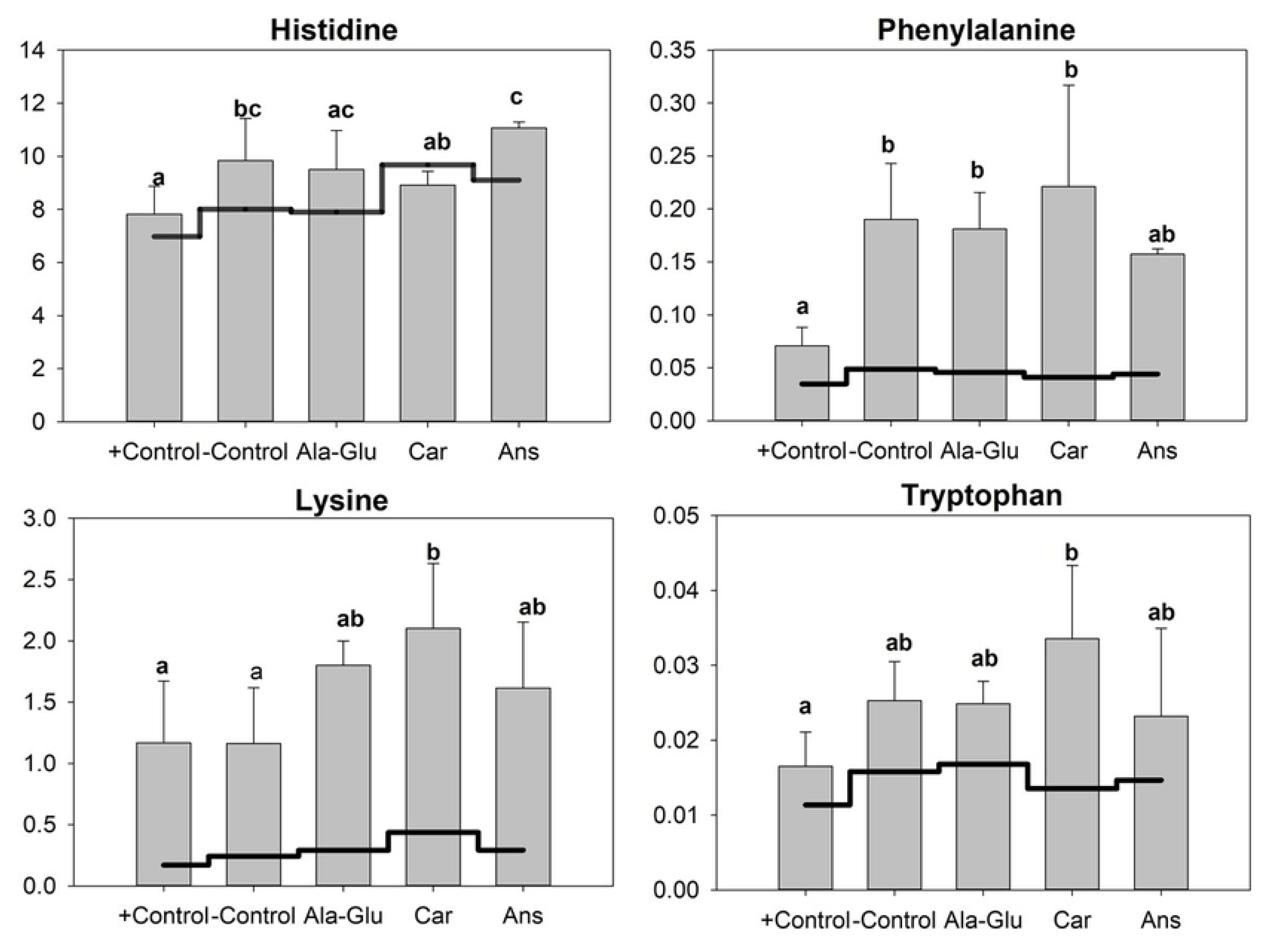
Indispensable free amino acid concentration. The concentration of indispensable free amino acids (µmol/g) in zebrafish muscle. Bars represent level (± std. dev) at 3hr after feeding. Line represents level at 24hr after feeding reflecting the baseline. Results of One-way ANOVA and LSD test are shown on graphs when significant. Significant differences (p<0.05) between groups are represented by different letters.

Among the dispensable free amino acids (DAA) the concentration of alanine was significantly lower in all groups compared to the (+) Control group (Figure 3). Tyrosine was significantly higher in all groups except Ans, compared to the (+) Control group. There were no significant differences in the concentration of arginine, except that Car had a significantly higher concentration than (+) Control. All groups had a significantly lower concentration of glycine compared to (+) Control (Figure 3).

**Fig 3.**
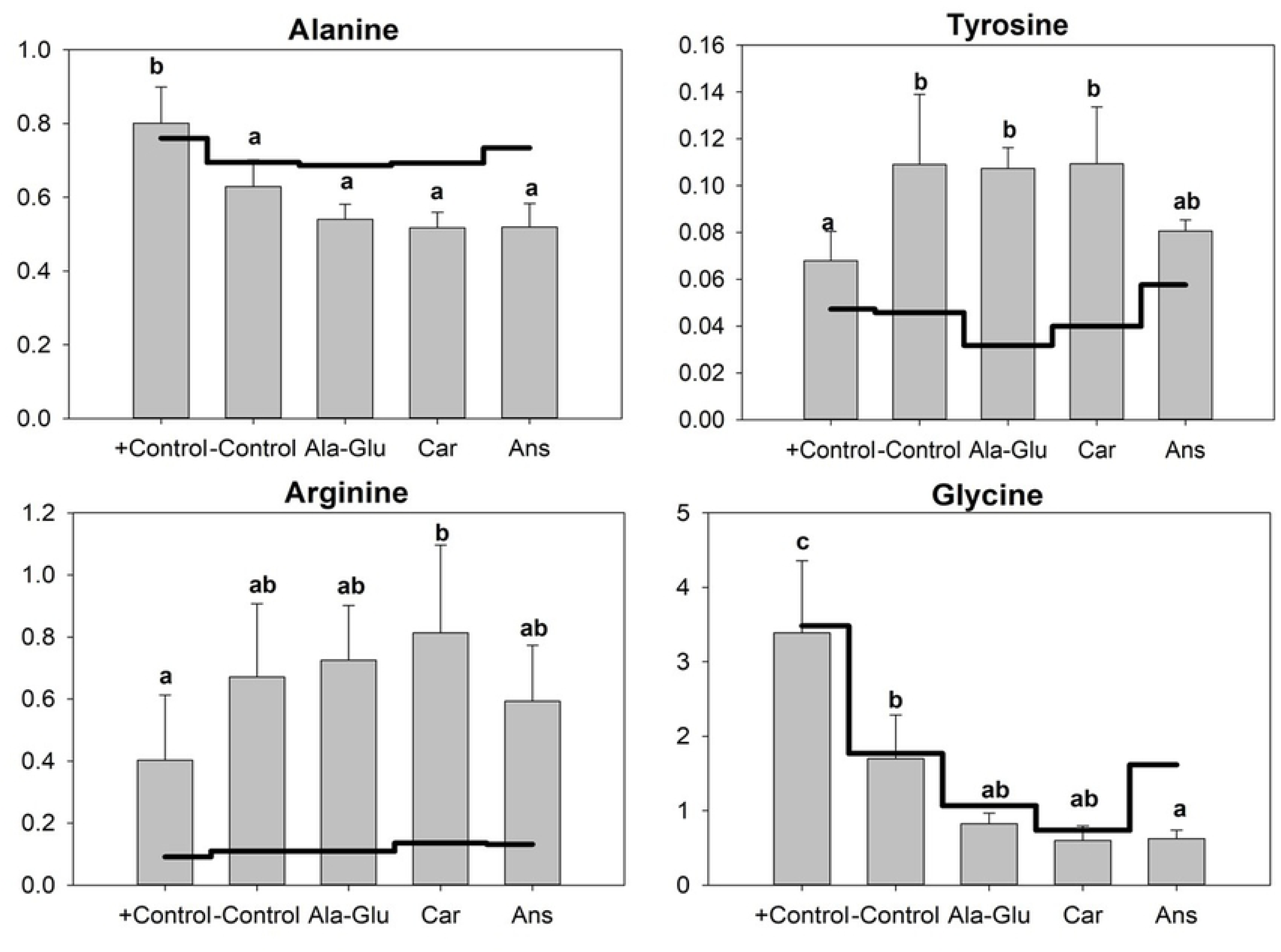
Dispensable free amino acid concentration. The concentration of dispensable free amino acids (µmol/g) in zebrafish muscle. Bars represent level (± std. dev) at 3hr after feeding. Line represents level at 24hr after feeding reflecting the baseline. Results of One-way ANOVA and LSD test are shown on graphs when significant. Significant differences (p<0.05) between groups are represented by different letters.

There were no significant differences in the levels of total IDAA or total free amino acids (TFAA) 3-hours after feeding among any of the groups. The (-) Control and Ans groups had significantly lower concentrations of DAA, compared to the (+) Control group but not different compared to Car or Ala-Glu groups (Table 5).

## Discussion

Previous studies have found that high dietary inclusion of PP significantly decreases the growth performance of many fish species [2,5,33]. Sitja-Bobadilla et al. [33] found that a 100% plant-based diet significantly reduced the specific growth rate (SGR) and feed efficiency of gilthead sea bream (*Sparus aurata*). In Atlantic Cod (*Gadus morhua*), PP inclusion levels above 50% reduced both the SGR and feed conversion ratio [2]. In addition to those results, our study found that the (-) Control group fed an SBM diet lacking dipeptide supplementation had a significantly reduced weight gain and average weight compared to the FM fed (+) Control group. However, the supplementation of SBM-based diets with alanyl-glutamine and carnosine significantly improved the growth performance of zebrafish. Compared to the (-) Control group, Ala-Glu and Car grew 35.38% and 33.96% larger, respectively. Other studies have looked at carnosine supplementation as a means of improving growth performance [15,34]. Snyder et al. [15], did not find that PP-based diets supplemented with carnosine significantly improved the growth performance of a carnivorous species, rainbow trout. Those authors, however, supplemented carnosine at a level of 0.39g/kg of diet, while this study used higher 1.0g/kg of carnosine in the diet. In addition, Snyder et al. [15] used diets based on soy protein concentrate and not SBM. The results from this study support the use of carnosine for improving the growth performance of fish fed a fully plant-based diet.

Focusing on intestinal health, previous studies have assessed the positive effects of carnosine supplementation on antioxidative capacity [34–36], and anti-inflammatory response [16,17,37]. Nagasawa et al. [36] determined that high concentrations of carnosine in muscle inhibited the oxidation of both lipids and proteins. Looking at the anti-inflammatory effects of carnosine, one pathway may be the inhibition of pro-inflammatory cytokines in the intestine [37,38]. The researchers found that expression of interleukin-8 in intestinal cells was down-regulated with the presence of carnosine in tissues [37]. Additionally, carnosine supplementation was able to significantly reduce expression of interleukin-6 and *tnf-α* in the liver of mice that were exposed to ethanol-induced inflammation [16]. However, supplementation of carnosine did not have a significant impact on the inflammatory cytokines that were assessed in this study.

The use of alanyl-glutamine supplementation as means of improving intestinal health started with positive results on mammals [19, 39–42]. Haque et al. [39] determined that the supplementation of alanyl-glutamine into the total parenteral nutrition of rats significantly improved functional morphology and metabolism within the intestine. This was observed through the increase of villus height, increase of mucosal protein, and a higher uptake of glutamine in the intestine [39]. In a mouse study, parenteral supplementation of alanyl-glutamine significantly increased both intestinal villus height and surface area [42]. Additionally, past research has looked into alanyl-glutamine supplementation and its effect on growth and intestinal health in fish [21,22,26,27]. In studies done on hybrid sturgeon (*Acipenser schrenckii*), dietary supplementation of alanyl-glutamine increased weight gain and growth rate [21,22]. These authors concluded that the increase in growth from the alanyl-glutamine supplemented group possibly resulted from improved intestinal health since the fish receiving supplementation showed significantly increased intestinal villus height and digestive enzyme activity [22]. Similar results were observed in this study; the Ala-Glu group showed longer intestinal villi than the (-) Control group and had a higher length-to-width ratio of intestinal villi. The intestinal villus length-to-width ratio provides an important measure of the efficiency of nutrient absorption within the intestine. A higher villus length-to-width ratio indicates increased surface area and ability for better absorption of nutrients [43], which correlates with an improved specific growth rate [44]. Reduced villus length-to-width ratio in the intestine has previously been found to be a result of intestinal inflammation caused by SBM, and has been associated with a decrease in the absorption capacity of enterocytes [8,9,44].

Nutrient absorption was assessed in this study by analyzing two genes, PepT1 and *fabp2*, measuring their level of expression within the intestine. PepT1 is a protein-bound transporter responsible for uptake of dietary di- and tripeptides [45]. PepT1 expression within the intestine was measured to determine if dipeptide supplementation stimulates PepT1 transport of those dipeptides, or if the studied dipeptides were being broken down into single amino acids prior to their absorption. Amino acids that are transported in peptide form are more readily absorbed for use in the body than amino acids transported in free form [46]. Expression of *fabp2*, a fatty-acid binding protein responsible for fat absorption within the intestine, was measured since past studies have found that SBM-induced intestinal inflammation significantly decreases expression of this gene [10, 44]. In this study, there were no significant differences in expression of either gene across the different experimental groups. An important trend that was observed however, was that the Ala-Glu group had numerically the highest expression of both genes, compared the other groups. Alanyl-glutamine supplementation has been previously observed to increase the expression of Pept1 [47,48]. Another study [41], found that alanyl-glutamine supplementation increased expression of PepT1 after 5 days after a small bowel resection. The increase in PepT1 could possibly signify that the dietary alanyl-glutamine is being absorbed by the intestine as an intact dipeptide [49,50]. While studies have not found a direct link between alanyl-glutamine supplementation and *fabp2* expression, the numerical increase in *fabp2* expression in the Ala-Glu group may be a result of the significant increase in surface area of the intestinal villi, which is correlated with increased absorption ability [43,44]. The intestinal health results of this study are consistent with a growth study done by Zhou et al. [48] on pigs, which found that the dietary supplementation of alanyl-glutamine improved intestinal health status and absorption capacity, resulting in a significant increase in growth of the studied animals. In fish, the alanyl-glutamine supplementation promoted anti-oxidation and in turn, improved the feed utilization and increased growth in juvenile cobia (*Rachycentron canadum*) [26]. The diet used in that study was only a partial PP feed, and 2/3 of the protein in the feed was FM based. Vitamin E supplementation was also tested in combination with alanyl-glutamine supplementation and the best results occurred in fish that were supplemented with both [26]. Our study builds on those results and provides support that alanyl-glutamine supplementation alone can increase the weight gain and total weight of fish that are fed a fully (100%) plant-based diet.

The replacement of FM with plant-based protein sources has been found to affect the dietary amino acid absorption in fish [2,51]. Previous research has found that inclusion of certain plant proteins in feed, can result in imbalanced FAA levels [52,53]. Mente et al. [52] reported that the inclusion of wheat gluten in the diets for Atlantic Salmon significantly reduced IDAA levels, especially threonine and lysine. Gomez-Requeni et al. [53] determined that the feeding of PP led to a decrease in arginine and lysine, and overall a lower utilization rate of dietary amino acids, which may have contributed to the decreased growth rate in the studied fish. In order to make up for the FAA imbalances that can be caused by using amino acid deficient PP sources, the amino acids in a crystalline form must be supplemented in the feed. However, FAA supplementation has been found to cause a saturation of brush border transporters, their catabolism and/or excretion, that eventually lead to a decrease in tissue protein synthesis [54,55]. Dietary amino acids supplemented in dipeptide form seem to improve growth performance and dietary amino acid utilization in salmonids [56,57]. The muscle FAA pool is key to understanding how the supplemented dipeptides affect dietary availability of certain amino acids and in this study especially alanine, glutamine, and histidine. The differences in concentrations of amino acids between the (-) Control group and the respective dipeptide supplemented groups were the focus to determine how each dipeptide affects the absorption and utilization of each amino acid within that dipeptide. The only FAA in muscle that was significantly different between the four groups fed SBM-based diet was lysine. Lysine concentration was higher in the Car group compared to the (-) Control group. Previous studies involving FM replacement with PP found lysine levels to be significantly reduced in the muscles of the fish [52,53]. Lysine is typically the first limiting amino acid when FM is fully replaced by PP [58]. The increased lysine level may indicate higher lysine availability and be one of the factors behind the increased growth performance of the Car group. On the contrary, a decrease in lysine level results in reduced protein retention and reduced growth performance [58,59]. It is possible that the increased level of free lysine in muscle in the carnosine-supplemented group may show that the improved growth performance is attributed to a higher digestibility of SBM, and hence increased retention of dietary amino acids within the muscle of the zebrafish. Carnosine supplementation has been reported to increase gastric secretion and digestive enzyme activity in rats [60]. Potential increase in digestive activity through carnosine supplementation may have improved the ability of the zebrafish to break down and absorb protein from the SBM feed better. This speculation, however, requires further investigation.

## Conclusion

The findings from this study are key to the improvement of utilization of plant-based feeds. The supplementation of alanyl-glutamine into an SBM-based diet significantly increased the growth performance of zebrafish and improved the surface area of the villi in the proximal intestine. Similarly, dietary carnosine supplementation improved growth performance, as well as increased the level of free lysine in the muscle. Both of these findings provide support for further research into alanyl-glutamine and carnosine additives as means of mitigating the negative effects of SBM-based feeds on growth performance in other fish species.

## Acknowledgements

We thank Saffron Scientific Histology Services (Carbondale, IL) for providing the histological services for this study.

